# Larval thermal characteristics of multiple ixodid ticks underlie their range dynamics

**DOI:** 10.1101/2019.12.11.873042

**Authors:** Alicia M. Fieler, Andrew J. Rosendale, David W. Farrow, Megan D. Dunlevy, Benjamin Davies, Kennan Oyen, Joshua B. Benoit

## Abstract

Temperature is a major factor that impacts tick populations by limiting geographic range of different species. Little is known about the thermal characteristics of these pests outside of a few studies on survival related to thermal tolerance. In this study, thermal tolerance limits, thermal preference, impact of temperature on metabolic rate, and temperature-activity dynamics were examined in larvae for six species of ixodid ticks. Tolerance of low temperatures ranged from −15 to −24°C with *Dermacentor andersoni* surviving at the lowest temperatures. High temperature survival ranged from 41 to 47 °C, with *Rhipicephalus sanguineus* having the highest upper lethal limit. *Ixodes scapularis* showed the lowest survival at both low and high temperatures. Thermal preference temperatures were tested from 0-41°C. *D. variabilis* exhibited a significant distribution of individuals in the lower temperatures, while the majority of other species gathered around 20-30°C. Activity was measured from 10-60°C, and the highest activity was observed in most species was near 30°C. Metabolic rate was the highest for most species around 40°C. Both activity and metabolic rate dropped dramatically at temperatures below 10°C and above 50°C. In summary, tick species vary greatly in their thermal characteristics, and our results will be critical to predict distribution of these ectoparasites with changing climates.

## Introduction

Ticks are blood-feeding, ectoparasitic arthropods that are known for spreading a wide range of diseases through very specific vector and host interactions (Wikel, 2013). Every life stage of ticks feed and often acquire diseases from vertebrate host when larvae feed (Riek et al., 1964). Lyme disease is one of the most commonly reported vector-borne illnesses (Stafford et al., 1998). Spotted fevers such as Rocky Mountain and *Rickettsia parkeri* are common diseases transmitted by the bacteria *Rickettsia* found in the saliva of the *Dermacentor variabilis, D. andersoni, Amblyomma maculatum* and *Rhipicephalus sanguineus ticks* (Parola et al., 2005). Tick-borne diseases have been consistently on the rise; recent studies have shown that the number of Lyme disease carrying ticks in Iowa has increased from 8% to 23.5% between 1998 and 2013 (Oliver et al., 2017) and the number of Lyme disease cases have been steadily increasing over time (Stafford et al., a1998). In addition, the occurrence rate of *Anaplasma phagocytophilum*, transmitted by *Ixodes scapularis*, and *Ehrlichia chaffeensis*, carried by *Amblyomma americanum*, has increased by more than 2-fold from 2000-2007 (Dahlgren et al., 2011). There are a multitude of putative reasons for this increased disease transmission, including improved recognition of diseases carried by ticks, climate change with shifts in the active season, distribution of ticks, and encroachment of humans into tick habitats (Lindgren et al., 2000).

The geographical distrubution of tick species is broadly known in the USA (Fig. 1); however, the current distribution is shifting, and its is expected that climate change will further alter the distribution of ticks in the future. Predicting future changes in tick distribution largely depends on the thermal characteristics displayed by each species. Ticks have demonstrated sensitivity to abiotic factors such as temperature and humidity, which is likely due to their long periods off host between blood meals (Needham et al, 1991; Rosendale et al., 2017; Springer et al., 2015; Yano et al., 1987; Yoder & Benoit 2003; Yoder et al., 1997, 2004).

**Figure 1:**
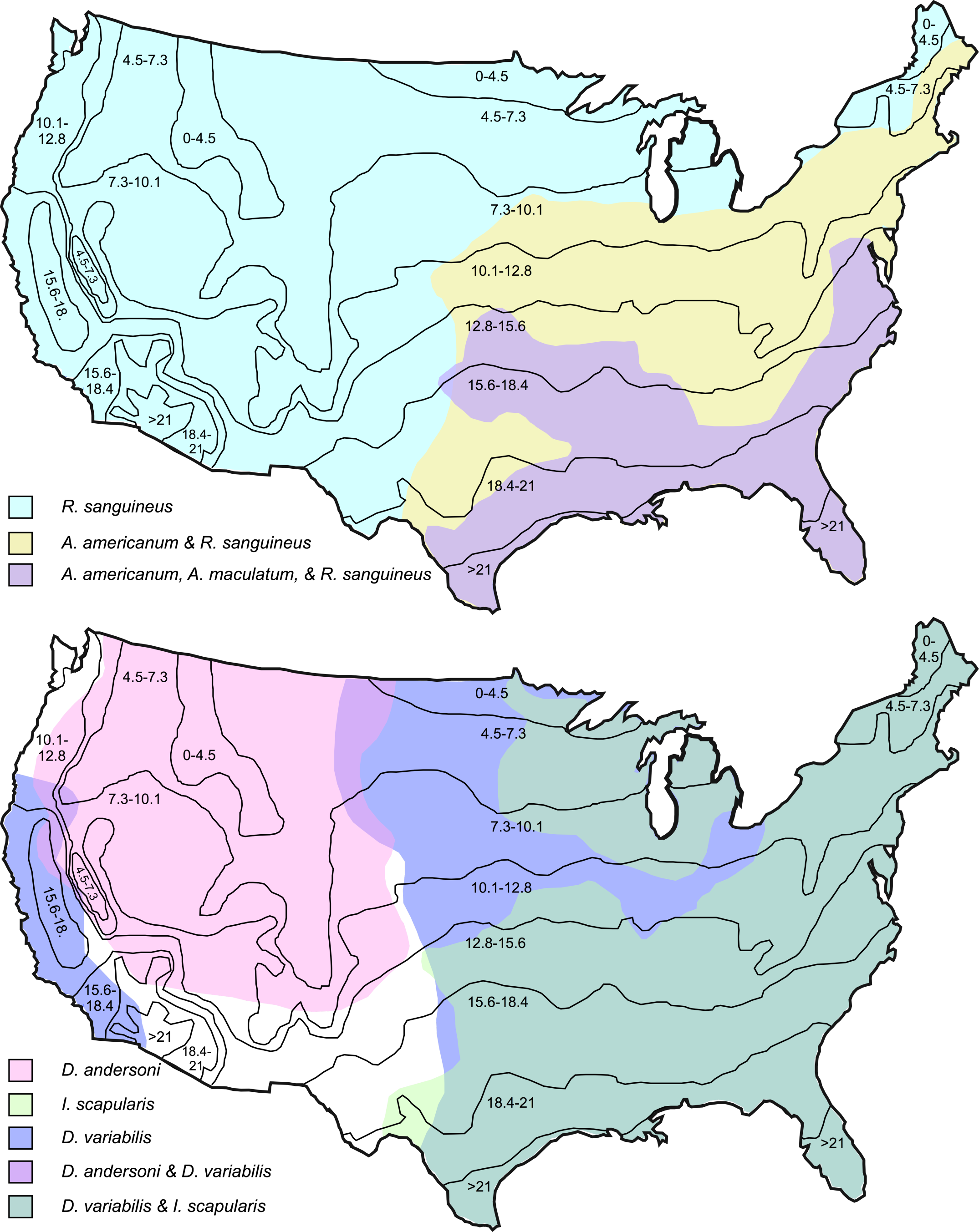
Geographic distribution of Ixodidae in the continental United States. Distribution data based on data available from the Centers for Disease Control and Prevention (CDC, 2018b).

Varying species of ticks are inept to surviving and acclimating to extreme high and low temperatures, where survival is impaired during short and long term exposure to temperatures that are too high and too low (Holmes et al., 2018). When exposed to such temperature extremes, ticks can become susceptible to dehydration and overheating in the summer, and freezing in the winter (Eisen et al., 2016; Rosendale et al., 2016; Yoder & Benoit 2003). Importantly, each bout of stress seems to yield a quantifiable expenditure of energy reserves, which cannot be replenished until a new host is located (Loye & Lane, 1988; Rosendale et al., 2017; Randolph et al., 1999). If temperatures become too high, lower questing activity in ticks has been noted (Loye & Lane 1988). This is likely since ticks will return to the duff layer of the soil to lower their temperate and rehydrate due to the higher relative humidity near the soil, which will decrease the probability of seeking a host (Tompkins, et al., 2014). Host-seeking behavior also declines with low temperature (Dautel et al., 2008); therefore, seasonal variations in temperature, and also photoperiod, determines activity patterns of ticks. These specific behavioral changes can ultimately impact the dynamics of tick-borne diseases by regulating host-tick interactions (Gilbert, 2010). Climate change also influences human behavior; a higher rate of individuals are spending more time outside, wearing fewer layers of clothing with subsequent higher exposure to ticks and potential diseases (Vázquez et al., 2008).

A consistent northern expansion of tick-borne diseases has been identified due to the increasing global temperatures, which poses an impact on tick dynamics (Gray et al., 2009). Studies on thermal characteristics of ticks have been lacking and solely focused on survival during exposure to lower and upper lethal temperatures without study of more physiological processes that could be impacted by thermal characteristics (Dautel & Knülle, 1997; Lee & Baust, 1987; Neelakanta et al., 2010). The purpose of this study is to examine thermal characteristics of ticks, including limits of thermal survival, effect of temperature on metabolic rate, activity patterns, and preferred temperature ranges of various tick species to predict future distribution patterns and the impact of vector-borne diseases. There are a limited amount of studies performed on tick thermal characteristics beyond survival studies; however, existing data indicates that global shifts in tick distribution correspond to the increasing climate change (Gray et al., 2018). Overall, our study demonstrates the influence that temperature has on the physiology and behavior of larval ticks. The most dramatic difference among species was seen at low temperatures, suggesting that winter conditions may have the greatest influence on tick behavior and distribution.

## Methods

### Ticks

Larvae of several species, including *Amblyomma americanum, A. maculatum, Rhipicephalus sanguineus, Ixodes scapularis, Dermacentor variabilis, D. andersoni*, were reared from eggs. Engorged females were acquired from laboratory colonies at the Oklahoma State University (OSU) Tick Rearing Facility (Stillwater, OK, USA). Adult ticks are maintained in these colonies at 14:10 h, light:dark (L:D), 97% relative humidity (RH), and 25±1°C. Mated females were fed on sheep (*Ovis aries*) until repletion. For our study, fed females were sent to our laboratory within several days of drop off from the host and, upon arrival, the engorged females were placed in closed chambers containing a supersaturated solution of potassium nitrate (93% RH; Winston and Bates, 1960). Females were allowed to lay eggs, and the eggs developed at 26±1°C, 15:9 h L:D, and 93% RH for several weeks. Multiple females (N=3) were used for each tick species to prevent results based upon a single clutch of eggs. After emerging, larvae were kept at these conditions until being used in experiments 2-4 weeks post-hatching.

### Thermal tolerance

Limits of thermal tolerance were determined based on studies of cold tolerance in *D. variabilis* (Rosendale et al., 2016). Briefly, larvae were subjected to high or low temperatures via 2 h exposures to cold (0 to −25°C) or heat (35 to 47°C).. Groups of 10 ticks (N= 10 groups) were placed in 1.5 cm^3^ tubes which were then arranged into foam-plugged 50 mL tubes, which were suspended in an ethylene glycol:water (60:40) solution. Temperature was regulated (± 0.1°C) with a programmable bath (Arctic A25; Thermo Scientific, Pittsburgh, PA, USA). After thermal treatments, larvae were returned to rearing conditions and allowed to recover for 24 h prior to survival assessment.

### Thermal preference

To determine the temperature preferred by the larval ticks, a thermal preference arena was constructed. The arena consisted on a 30 cm long piece of clear plastic tubing (2.5 cm diameter) that was heated at one end with water (40°C) circulated from a programmable bath and cooled on the other end with and ice bath. The whole length of tubing was placed within insulated boxes. This setup resulted in one end of the arena being at 43°C while the other end was at 0°C and the middle was ∼24°C. The arena was visibly marked along its length into multiple sections with known temperatures. Groups of 10 (N=10 groups) ticks were released into the center of the arena and allowed to freely move for 2 h. A total of 5 trials were performed and at the end of each trial, the number of ticks in each section was recorded.

### Oxygen Consumption

The effect of temperature on O_2_ consumption in larval ticks was examined using microrespirometers constructed and utilized as previously described (Lee, 1995; Rosendale et al., 2017). Groups of 10-20 larvae (n=10 groups) were positioned within the microrespirometers such that ∼0.02 cm^3^ of space was available for the ticks. Microrespirometers were positioned in heated or cooled baths (0 - 60°C) and the complete apparatus was allowed 15 minutes to equilibrate prior to measurements being taken. CO_2_ production, measured through the movement of KOH, was measured every hour for 6 hours. Assuming that one mole of O_2_ is consumed for every mole of CO_2_ that is released, oxygen consumption was calculated by using the distance travelled by the KOH and was expressed as nl O_2_ mg^-1^ wet mass tick^-1^ h^-1^.

### Activity

Activity was monitored using a Locomotor Activity Monitor from Trikinetics Inc. (Waltham, MA, USA) and the DAMSystem3 Data Collection Software (TriKinetics). Larva (*n* = 20) were individually placed in 4 inch glass test tubes which consisted of tightly packed cotton at the top and bottom so that the ticks were contained in a 12.5 cm^3^ space in the middle. The tubes were loaded horizontally into the locomotor activity monitor and the entire apparatus was placed inside an insulated plastic container that contained water to maintain a high relative humidity (>97%) throughout the trial. Water that was heated or cooled with a circulating water bath was circulated through plastic tubing within the insulated container to generate temperatures ranging from 10-60°C +/- 2°C. Activity was measured by a sensor that counted the number of times the tick crossed the center of the tube, this number is directly related to the activity and movement of the ticks.

### Statistical analyses

Summary statistics are reported as means ± standard error (SE). For survival data, the proportion of larvae that survived in each tube was recorded and all survival data were arcsin-square root transformed prior to analysis. Using Prism (GraphPad; San Diego, CA, USA), survival, activity, and oxygen consumption were analyzed using a two-way analysis of variance (ANOVA) with species and temperature as the factors, and pairs of means were compared among species using Tukey’s *post-hoc* tests. For thermal preference, means were compared using a one-way ANOVA with a Tukey’s *post-hoc*.

## Results

### Thermal tolerance

The lower lethal temperature (LLT), where no larvae survived a 2 h exposure, ranged among species from −15 to −24°C (Fig 2, Table 1) with *A. americanum* and *I. scapularis* having the highest LLT and *D. andersoni* having the lowest. The temperature resulting in 50% mortality (LLT_50_) varied (*F*_5,43_ = 175, *P* < 0.0001) among species with *R. sanguineus* and *I. scapularis* having the highest LLT_50_ and *D. andersoni* having the lowest (Table 1). There was a significant effect of temperature (*F*_10, 603_ = 379.4, *P* < 0.0001), species (*F*_5, 603_ = 231.2, *P* < 0.0001), and a significant temperature*species interaction (*F*_50, 603_ = 19.7, *P* < 0.0001). Overall survival varied among all species (*P* < 0.01, all cases) with the exception of *R. sanguineus* and *I. scapularis*, which did not differ from each other (*P* = 1.0).

**Table 1.** Thermal tolerance of larval ixodid ticks.

|  | LLT | LLT<sub>50</sub> | ULT | ULT<sub>50</sub> |
| --- | --- | --- | --- | --- |
| *A. americanum* | -15 | -11.4 ± 0.5<sup>a</sup> | 45 | 40.5 ± 0.2<sup>a</sup> |
| *A. maculatum* | -22 | -12.1 ± 0.3<sup>a</sup> | 46 | 42.3 ± 0.3<sup>b</sup> |
| *D. andersoni* | -24 | -19.7 ± 0.3<sup>b</sup> | 45 | 41.1 ± 0.2<sup>a</sup> |
| *D. variabilis* | -20 | -16.9 ± 0.4<sup>c</sup> | 43 | 40.2 ± 0.2<sup>a</sup> |
| *I. scapularis* | -15 | -8.5 ± 0.3<sup>d</sup> | 41 | 38.0 ± 0.2<sup>c</sup> |
| *R. sanguineus* | -16 | -7.9 ± 0.3<sup>d</sup> | 47 | 42.3 ± 0.3<sup>b</sup> |

Values (mean ± SEM) that do not share a letter are significantly different.

**Figure 2:**
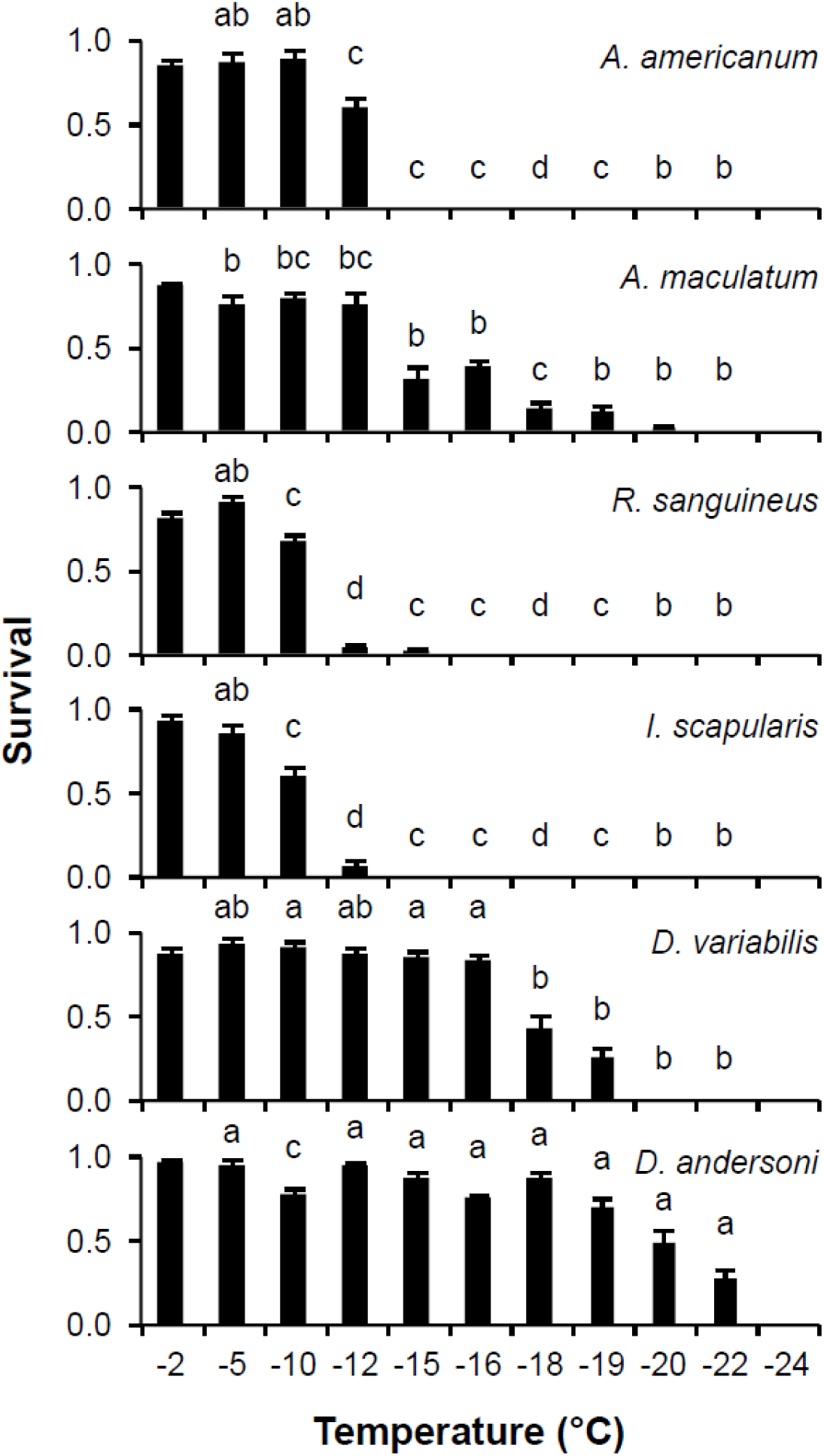
Survival of ixodid larvae exposed to thermal stress. Tick larvae were exposed to low temperatures for 2 h. For each temperature, values (mean ± SE) that do not share a letter are significantly different among species.

The upper lethal temperature (ULT), where no larvae survived a 2 h exposure, ranged among species from 41 to 47°C (Fig 3, Table #) with *I. scapularis* having the lowest ULT and *R. sanguineus* having the highest. The temperature resulting in 50% mortality (ULT_50_) varied (*F*_5,35_ = 35.0, *P* < 0.0001) among species with *I. scapularis* having the lowest ULT_50_ and *R. sanguineus* and *A. maculatum* having the highest (Table #). There was a significant effect of temperature (*F*_8, 425_ = 229.2, *P* < 0.0001), species (*F*_5, 425_ = 44.9, *P* < 0.0001), and a significant temperature*species interaction (*F*_40, 425_ = 9.0, *P* < 0.0001). Overall survival varied among the species with *I. scapularis* having the lowest survival and *R. sanguineus, A. maculatum*, and *D. andersoni* having the highest.

**Figure 3:**
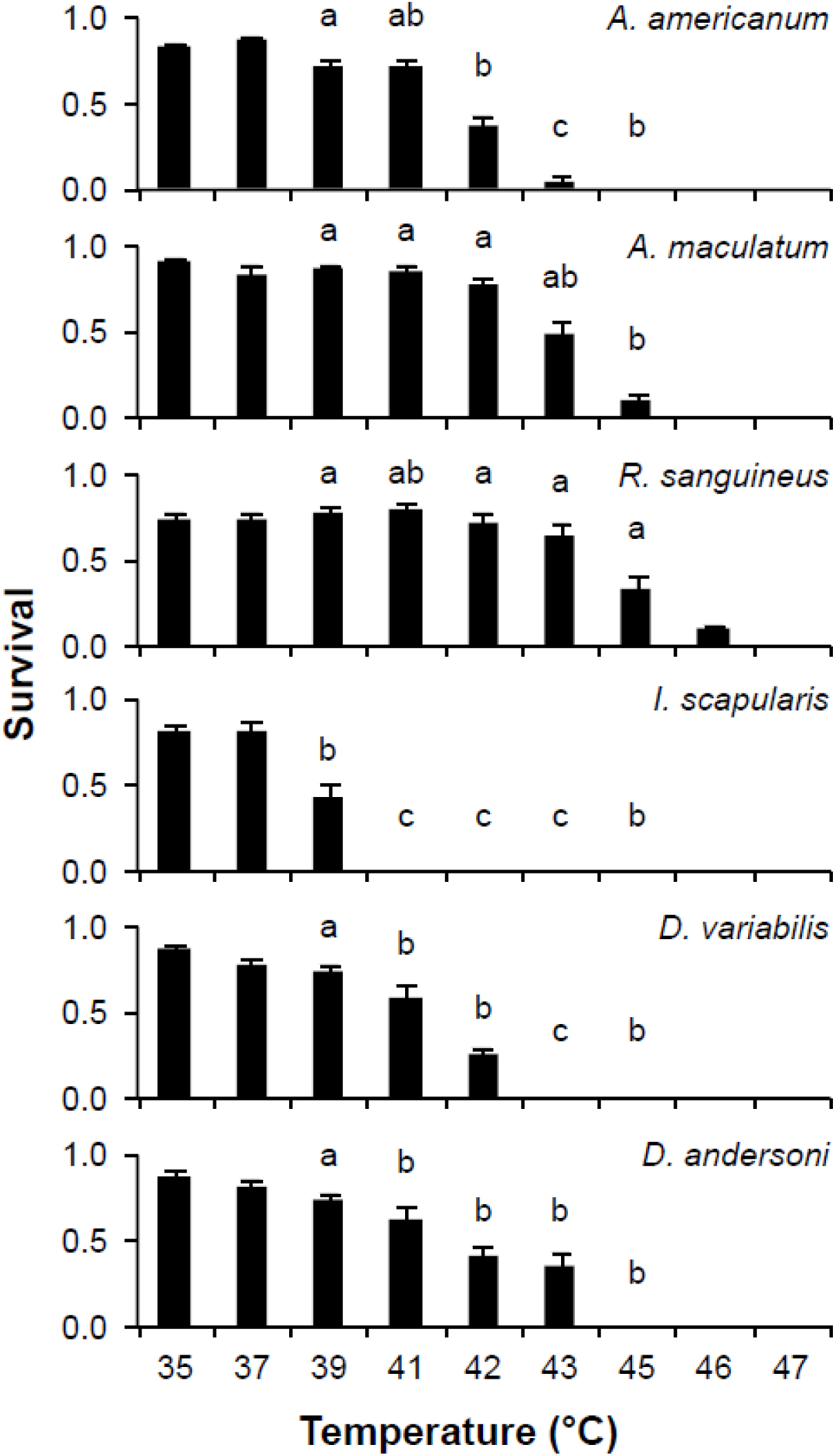
Survival of ixodid larvae exposed to thermal stress. Tick larvae were exposed to high temperatures for 2 h. For each temperature, values (mean ± SE) that do not share a letter are significantly different among species.

### Activity

To examine the effects of temperature on tick activity, larvae were placed in horizontally positioned tubes at temperatures ranging from 10 to 60°C, and their activity (number of times they moved across the center of the tube) was monitored for ∼24 h. Activity was highest at 30°C for most species, with *D. variabilis* and *A. americanum* having highest activity at 40°C (Fig. 4). Activity was essentially reduced for all species at 10 and 60°C. The lack of activity at 50 and 60°C is likely a result tick death. There was a significant effect of temperature (*F*_6, 598_ = 44.7, *P* < 0.0001), species (*F*_5, 598_ = 4.3, *P* = 0.0007), and a significant temperature*species interaction (*F*_30, 598_ = 6.2, *P* < 0.0001). Variation in activity levels occurred among several species at 20, 30, and 40°C.

**Figure 4:**
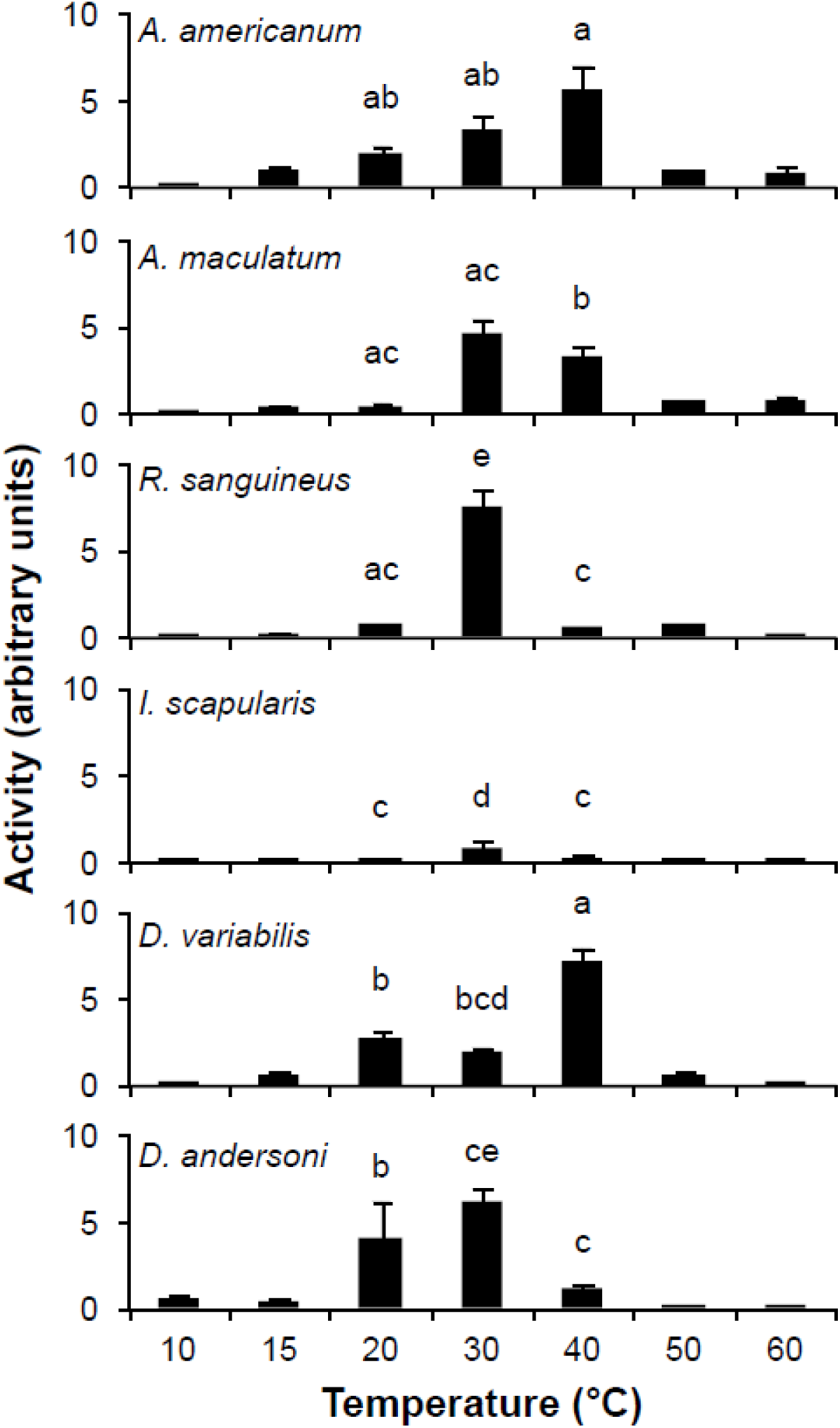
Activity of ixodid larvae at various temperatures. Activity levels were monitored for 24 h periods. For each temperature, values (mean ± SE) that do not share a letter are significantly different among species.

### Respirometery

The effect of temperature on oxygen consumption rates was examined by placing microrespirometers containing groups of larvae at various temperatures. Metabolic rate was highest at 40°C for most species, with *I. scapularis* and *A. maculatum* having highest rates at 30°C (Fig. 5). Oxygen consumption stopped and/or dropped below our ability to measure for all species at 0°C. At the upper thermal temperature, a drastic reduction in oxygen consumption occurred at 50°C and on differences were detected at 60°C (Fig. 5). This rapid decline is likely due to morality of ticks during prolonged exposure to high temperatures. There was a significant effect of temperature (*F*_5, 175_ = 149.2, *P* < 0.0001), species (*F*_5, 175_ = 13.3, *P* < 0.0001), and a significant temperature*species interaction (*F*_25, 175_ = 5.1, *P* < 0.0001). Variation in metabolic rate occurred among species at 30 and 40°C.

**Figure 5:**
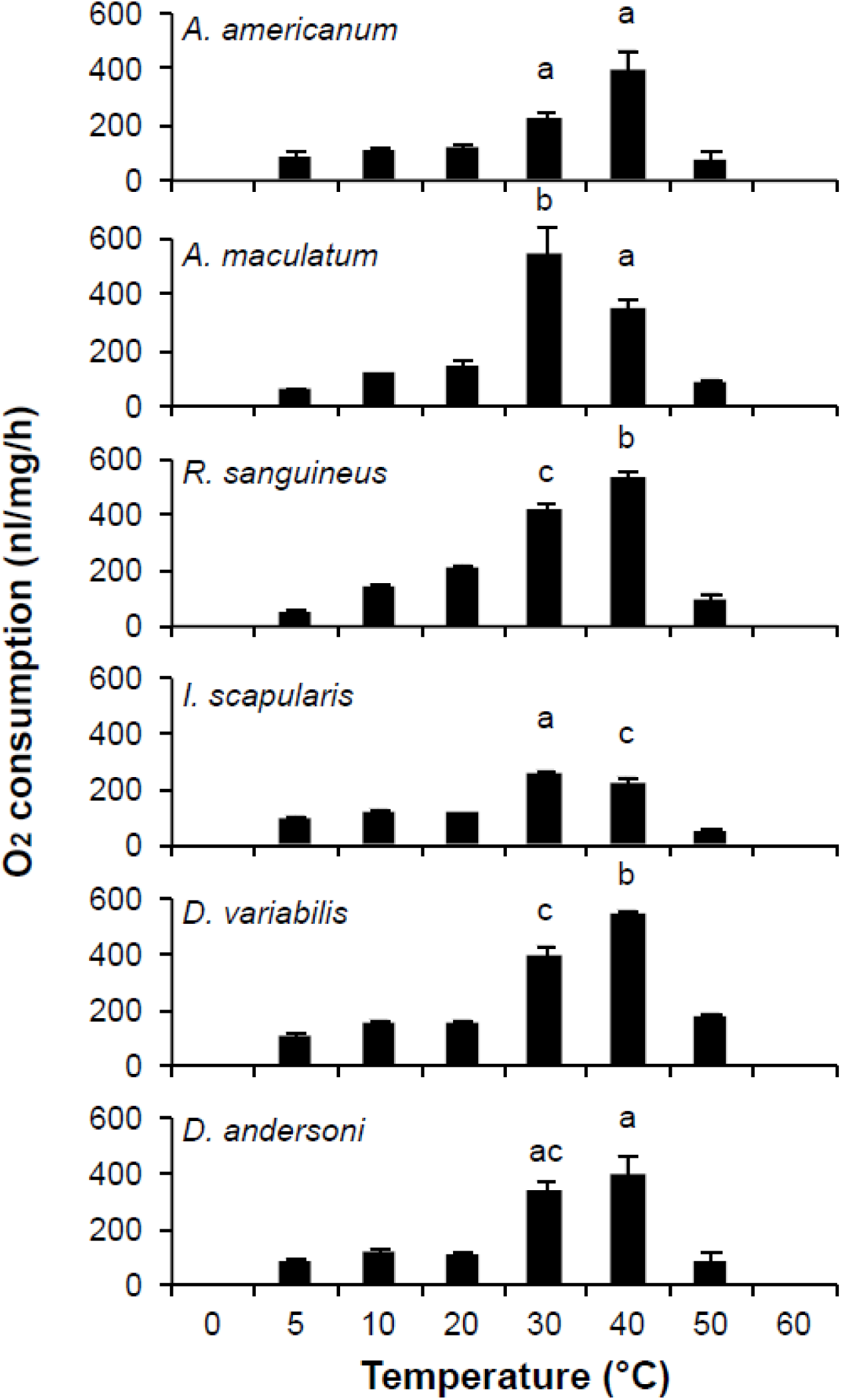
Effect of temperature on oxygen consumption of ixodid ticks exposed to various temperatures. Metabolic rate was monitored for 6 h. For each temperature, values (mean ± SE) that do not share a letter are significantly different among species..

### Thermal preference

To determine the preferred temperature of the different tick species, larvae were positioned in the center of a length of tubing that was cooled (0°C) at one end and heated (41°C) at the other and allowed 2 h to move to their preferred location. Although there was some variation among species as to which temperature had the highest proportion of individuals, most larvae aggregated in the 20-30°C temperature range (Fig. 6). The notable exception was *D. variabilis*, which had a higher proportion of individuals at the low temperature range. When the total distribution of the individuals was considered, there was significant (F5, 273 = 10.9, P < 0.0001) variation among the species, with *D. variabilis* being lower (P < 0.05, all cases) than all other species (Fig. 6).

**Figure 6.**
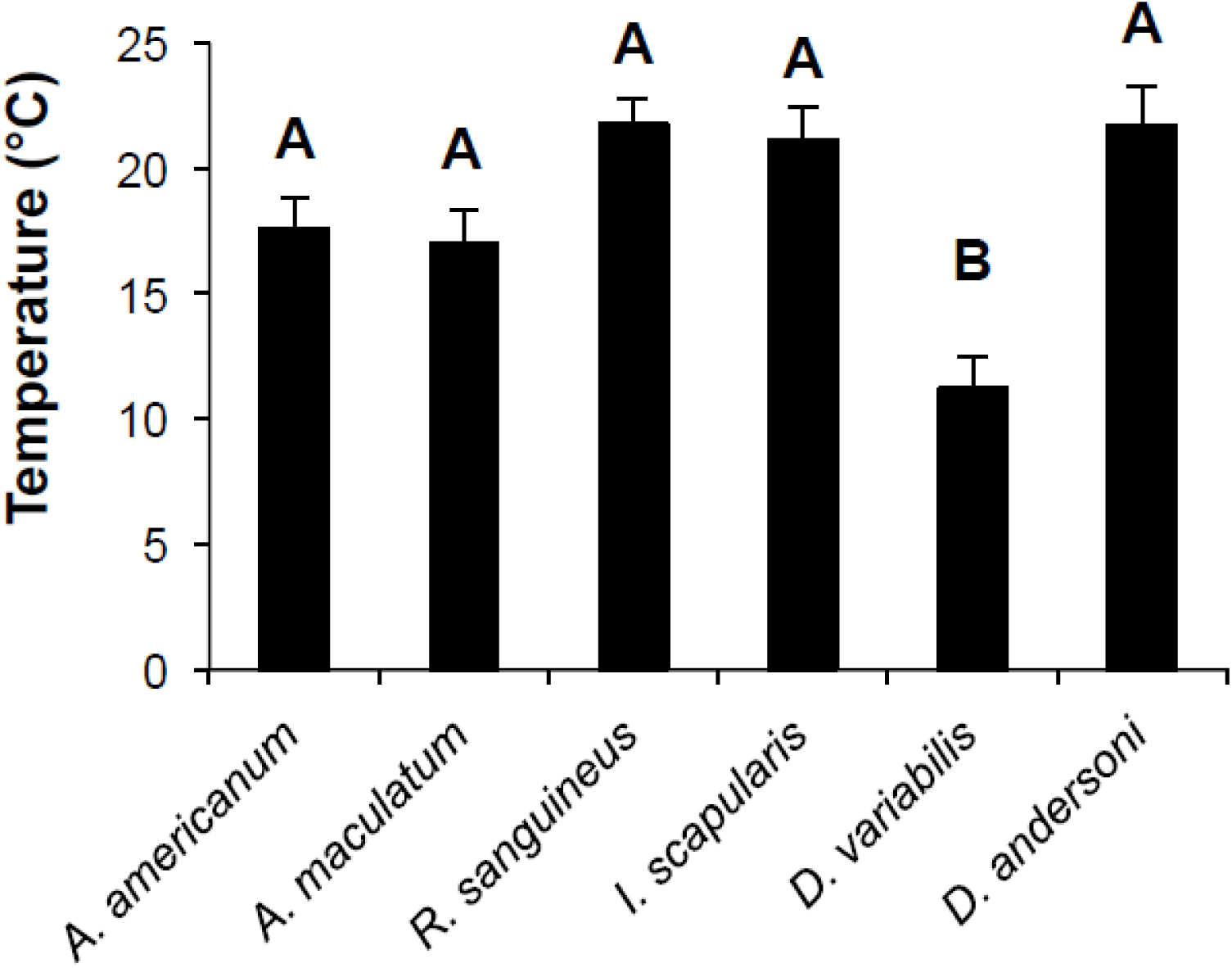
Thermal preference of ixodid ticks. Ticks were subjected to temperatures ranging from 0-41°C for 2 hrs. For each temperature, values (mean ± SE) that do not share a letter are significantly different among species.

## Discussion

The capability of ticks to survive in unfavorable environmental conditions contributes to their success as vectors of disease. Temperature fluctuations and other bouts of stress have been found to affect tick questing behaviors, metabolic rate, and energy reserves (Gilbert et al., 2014; Rosendale et al., 2016a; Rosendale et al 2017; Randolph et al., 1999). Thermal tolerance was examined in ixodid ticks by determining the lower lethal temperature (LLT) and the upper lethal temperature (ULT) after exposure larva to a wide range of temperatures, allowing for comparisons among several different species. The majority of species had a high survivability at −12 °C with survival decreasing dramatically after −16 °C. *D. andersoni* exhibited the highest tolerance to the low temperatures at ∼ 24°C whereas *A. americanum* and *I. scapularis* exhibited the lowest at ∼ −12°C. This supports the knowledge that *D. andersoni* is primarily located in the colder northern regions and higher elevations of the Rocky Mountains and can exhibit the adaptations needed to survive in lower temperatures (James et al., 2006). Rosendale et al. (2016) previously described these cold adaptations contributing to the success of cold tolerance including reduced photophase, dehydration and long term thermal acclimation. When examining higher temperatures, *R. sanguineus* exhibited the highest upper lethal temperature of approximately 47°C. and *I. scapularis* exhibited the lowest at 41°C. Results of the upper thermal limits are similar to those of prior studies described by Yoder et al. (2009) that displayed *R. sanguineus* having a high heat tolerance. *Amblyomma* species which are primarily located on the South Eastern coast, were described as being relatively heat tolerant which is supported by this study and previous research on survival (Yoder et al., 2009). Tolerance of upper lethal temperatures is not widely studied, but prior research suggests heat shock and stress response proteins contribute to their success in surviving higher thermal temperatures (Villar et al., 2010). The results of the higher temperatures may not be considered a major factor limiting temperature range as high temperatures had much less variation than the cold temperatures in relation to survival. The ranges tolerated were significantly more wide-spread at the lower temperature range, indicating that there may be more physiological differences among species related to low temperature survival that may play a role in determining thermal tolerance.

Ticks are hardy and can sustain survival for over a year between blood feedings (Lighton and Fielden, 1995; Rosendale et al., 2018). Increased temperatures have been correlated with an increase in metabolic rate as the kinetic processes that influence the biochemical reactions speed up (Addo et al., 2002). This increase in metabolism is also directly correlated to oxygen consumption and is an important factor in regulating chemical reactions needed to maintain homeostasis (Hawkins et al., 1995). To examine metabolism among several species of ticks, oxygen consumption was tested. A controversial topic of ectotherms is metabolic cold adaptation, which is a physiological adaptation of elevated basal metabolic rate to cope with temperature fluctuations (Lee et al., 1995). Ticks have a standard metabolic rate that is lower than other arthropods due to a low ratio of actively respiring tissue in relation to their body (Lighton et al., 1995; Rosendale et al., 2018). Metabolic rate was evaluated with all species displaying a peak O_2_ consumption rate between 30°C and 40°C. This peak in O_2_ consumption supports a relationship between increased metabolic rates with rising temperatures to the point where increased temperature (Pörtner et al., 2000). Above 40°C, metabolic rate decreased significantly, likely due decreases in performance above larvae due to thermal stress. These thermal performance curves can be used as much better indicators, rather than single measurements, for the predictions of organismal survival with changing climate change (Sinclair et al., 2016 along with Sinclair 2012 and Tuzun and Stock 2018).

Dynamics between temperature and metabolic rates of ticks will influence tick behaviors such as activity. The temperature range of 30°C and 40°C displayed the peak in activity levels, which corresponded with the peak in oxygen consumption. This peak in activity is likely an attempt to find an area with more optimal temperatures to reduce energy expenditure (Hawkins et al., 1995). A wide variety of invertebrates display strong temperature preferences due to specific thermoreceptors (Hamada et al., 2008). Preferential thermoregulatory behaviors have linked to optimal performance levels, where a delicate balance between preventing the rapid decline of metabolic reserves and locating a new host must be balanced (Abram et al., 2017). Preferential temperature ranges of ticks was between 15°C and 25°C with the exception of *D. variabilis*, which displayed a higher proportion of ticks at 12°C. The lowered temperature preference in *D. variabilis* could be a higher tolerance to colder temperatures (Rosendale et al., 2016b; Yu et al., 2014) or an impaired ability to detect temperatures. Importantly, is unlikely that *D. variabilis* was immobilized by the lower temperature since activity was noted as low as 10°C.

Overall, this study demonstrates a relationship between thermal characteristics of ticks and historic geographical distribution. The combined study of activity, oxygen consumption, and thermal indicates that the optimal performance of most ticks is likely between 20-40°C. Species near the lower end of the spectrum, such as *D. variabilis* and *D. andersoni*, will likely have reduced distributions as temperatures warm and vice-versa for warm adapted species, *A. americanum, A. maculatum*, and *R. sanguineus*, These trends are supported by our survival data, which have similar trends. Tick thermal characteristics has been scarcely studied in ticks, and a better understanding of these cold-related physiological shifts will allow us to further examine tick physiology as well as predict and identify future distribution and disease patterns.

## Acknowledgments

This work was supported by the University of Cincinnati Faculty Development Research Grant to J.B.B. Funding was also provided by the United States Department of Agriculture National Institute of food and agricultureGrant 2016-67012-24652 to A.J.R. Partially funding was to JBB was provided by the National Science Foundation DEB-1654417.

